# A preliminary study on the cistrome of human postsynaptic density from an evolutionary and network-based perspective

**DOI:** 10.1101/2021.01.25.428072

**Authors:** Zsofia E. Kalman, Zoltán Gáspári

## Abstract

The postsynaptic density (PSD) is a neuronal organelle that consists of thousands of protein complexes, having a role in signal transduction. The emergence of the complexes is dependent on the presence of proteins provided by gene expression. In this research we used Chip-seq data supported by protein level information. We developed a pipeline using data from five neuronal transcription factors, which reduces the false-positive hits of identified binding sites. In addition we found correlation between co-regulation and protein complex formation. The developed method paves the way for a future for large scale analysis utilizing a more comprehensive set of transcription factors.

## Introduction

The postsynaptic density (PSD) is a specialized region located below the membrane of the dendritic spine of a postsynaptic neuron. Its primary function is to receive transmitters released from presynaptic neurons and transmit electrical and biochemical signals in the postsynaptic cell [1]. PSD proteins can be divided into different structural and functional groups: membrane proteins and scaffold proteins are structurally dominant, while functionally cytoskeletal and regulatory proteins (phosphatases, kinases, and other regulators) are the most prevalent classes [1]. PSD plays a key role in shaping synaptic plasticity, which is essential for memory and learning [2]. A popular theory gaining ground lately is that the born and learned behaviour patterns are controlled by a diverse population of proteins that form a highly integrated and complex network [1]. The exact composition of this dynamically changing protein-protein interaction (PPI) network is influenced by several processes, ranging from protein expression to metabolic changes. Complex assembly is also governed by the structural composition of PSD proteins, including many domain-domain or domain-motif interactions [3]. The importance of the PSD is further enhanced by the fact that perturbation of certain hub proteins in the network has already been associated with more than hundred genetic diseases [4]. Unfortunately our current knowledge is largely limited to the pairwise interactions and to smaller complexes.

The evolution of PSD has already been studied at the protein and genomic levels, with several processes already identified (e.g., genome rearrangements) to contribute to the development of synaptic complexity [5]. However, it may also be interesting to investigate non-coding regulatory elements (Transcription Factor Binding Site, TFBS) from an evolutionary aspect. Variations in non-coding regulatory elements can lead to alterations in the abundance of PSD proteins and thus to phenotypic differences. Furthermore, in the case of many diseases mutations affecting the regulatory regions can lead to changes that will eventually manifest at the protein domain level as well [6,7].

In this work, Chip-seq data was analyzed from two perspectives: we investigated evolutionary changes of TFBSs, which may have an effect on the increase in nervous system complexity. Furthermore, to assess how these differences influence the composition of the PPI network of PSD, we examined the occurrence of genes responsible for PSD proteins near to these co-regulatory patterns.

## Methods

The neuronal Chip-seq data set was derived from the Chip-Atlas [8], and after manual verification, BED and NarrowPeak format data were downloaded from the NCBI-GEO [9]. Hg19 sequence data were obtained from the UCSC[10]. The synaptic gene data set was derived from SynaptomeDB [11]. Frequency matrices of selected transcription factors were downloaded from the JASPAR [12] database. The emf files for the multiple sequence alignments were downloaded from the Ensembl [13] FTP server. CHEA3 was used to obtain co-regulatory data [14], and Intact for protein interactions [15]. The FIMO (Find Individual Motif Occurrence) method was used to determine transcription binding sites [16].

## Results and discussion

### Chromosomal distribution of TFs is broadly similar except for NRF1

In our study we investigated the binding sites of five transcription factors (Gata2, Hand2, Jund, Nrf1, and Phox2b), previously indicated to play important roles in neuronal cells (e.g., regulation of differentiation, regulation of proteins involved in ubiquitination) [17][18][19][20][20,21]. Aligning Chip-seq experiment results to the reference genome results in peaks with variable intensity, containing several binding sites. We downloaded human neuronal sequences from NCBI GEO and used the FIMO server with default settings to identify the transcription factor binding sites (TFBSs). Table 1 shows the identified binding sites using different frequency matrices.

**Table 1:** Examined transcription factors, the number of bindings sites identified and the frequency matrices from JASPAR.

| Transcription factor | Identified sites with FIMO | Conserved identified sites | % of conserved identified sites | Number of sequence logos from Jaspasr |
| --- | --- | --- | --- | --- |
| GATA2 | 37 684 | 9 935 | 26.26 % | 3 |
| NRF1 | 6 246 | 331 | 5.2 % | 1 |
| HAND2 | 21 945 | 13 559 | 61.79% | 1 |
| JUND | 41 551 | 11 979 | 28.83% | 2 |
| PHOX2B | 156 816 | 83 898 | 53.50% | 1 |

To overcome the usually high false-positive rate of identified TFBSs, we also investigated the chromosomal distributions. We expect that true hits will more likely show a unique pattern, while similar occurrences may indicate general bias. In this case, we found that, with the exception of NRF1, TBFS showed a broadly similar distribution (Figure 1).

**Figure 1.**
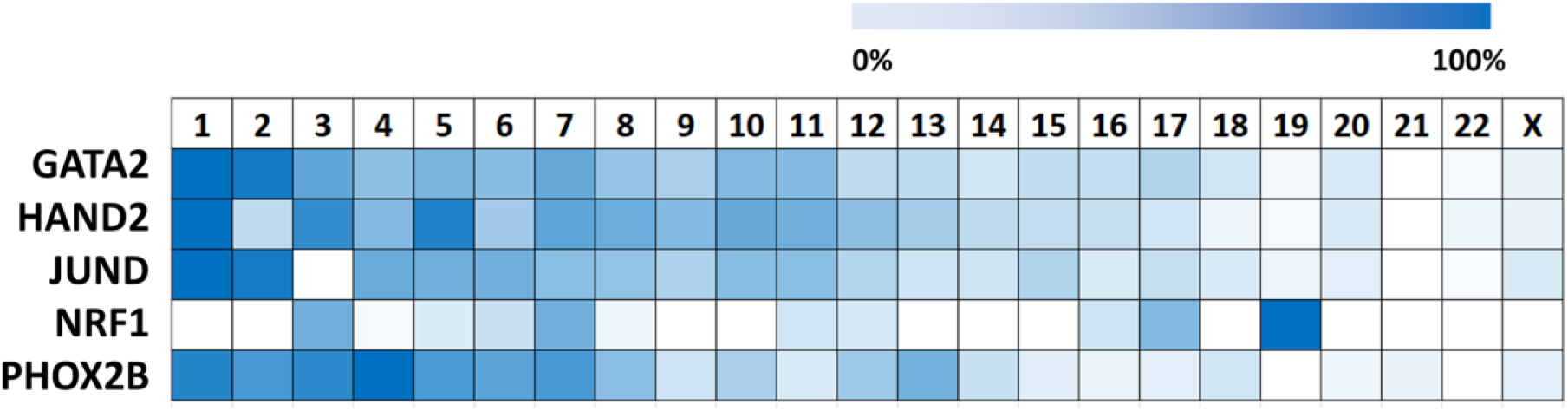
Distribution of transcription factors in different chromosomes.

To investigate the evolutionary differences between species we also downloaded the corresponding multiple sequence alignment from Ensembl and identified the regions belonging to each TFBSs. The multiple alignments contained 36 vertebrate species from rodents to primates. In general the quality of the alignments for the non-coding regions are lower compared to the protein-coding portions. Therefore as a result, we were unable to map all the binding sites back to the human genomic positions. The number of identified TFBS for each TFs that show conservation is shown in Table 1. There are considerable differences between the proportions of successfully aligned TFBSs. For PHOX2B we were able to map more than 50% of the TFBS to the multiple sequence alignment, while in the case of NRF1 it was around 5% - probably corresponding to high sensitivity/low specificity for the former one and vice versa for the latter one (Table 1).

### Quality of the identified binding sites

As a further analysis of the data, we examined the coverage of the motifs in the multiple sequence alignments (for 36 species). Figure 2 shows how the identified TFBSs are actually present in the position as FIMO indicated across various species. Coverage appears to be broadly adequate and in the vast majority of cases it was quite high.

**Figure 2:**
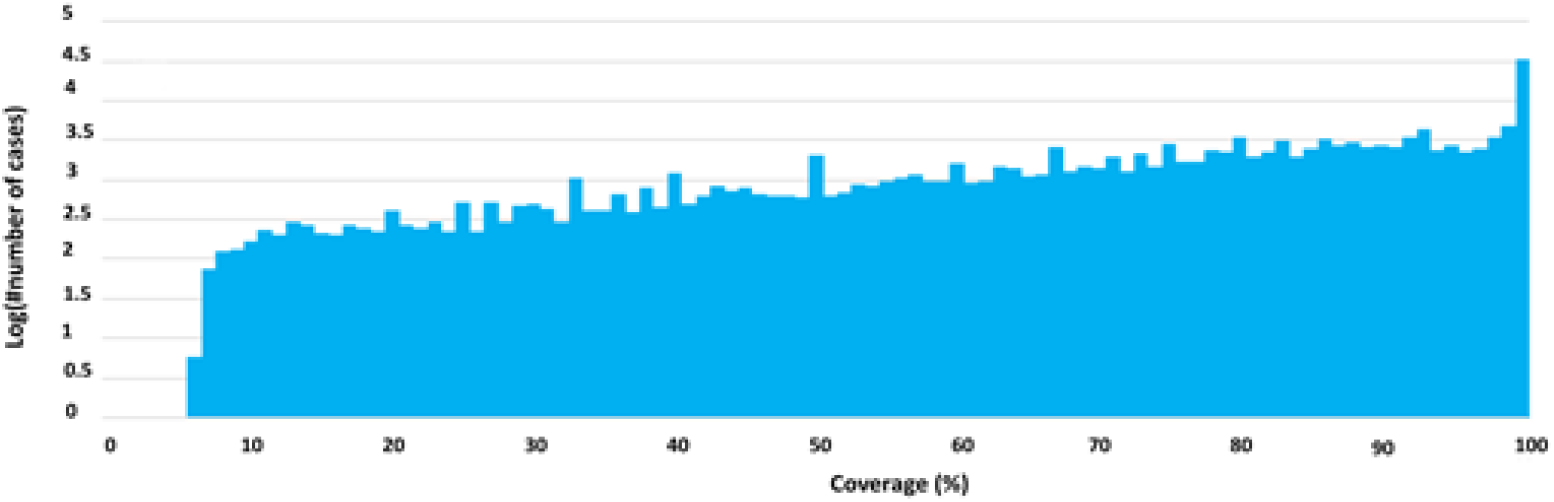
Distribution of coverage of all motifs across different species. Y-axis shows the log(number of motifs), while X-axis shows the coverage.

We also compared the sequence logos, the corresponding frequency matrices downloaded from Jaspar and the identified human TBFSs. We calculated the probability for each motif’s expected abundance using the frequency matrices. The sequence logos of JUND, GATA2, PHOX2B and HAND2 are less specific, resulting in lower probability values and very likely detection of a large number of false-positives, in contrast to the highly specific NRF1 logo, where probabilities are much higher (Figure 3). The higher specificity observed for NRF1 are on par with lower number of (conserved) hits, allowing much less space for sequence variability in the aligned region (see Table 1). FIMO probability values do not differ highly in case of NRF1 compared to other TFs, advocating the necessity of the utilized quality controls. Notably, sequence logos have limitations in terms of descriptions, as they cannot handle logical relationships and gaps.

**Figure 3:**
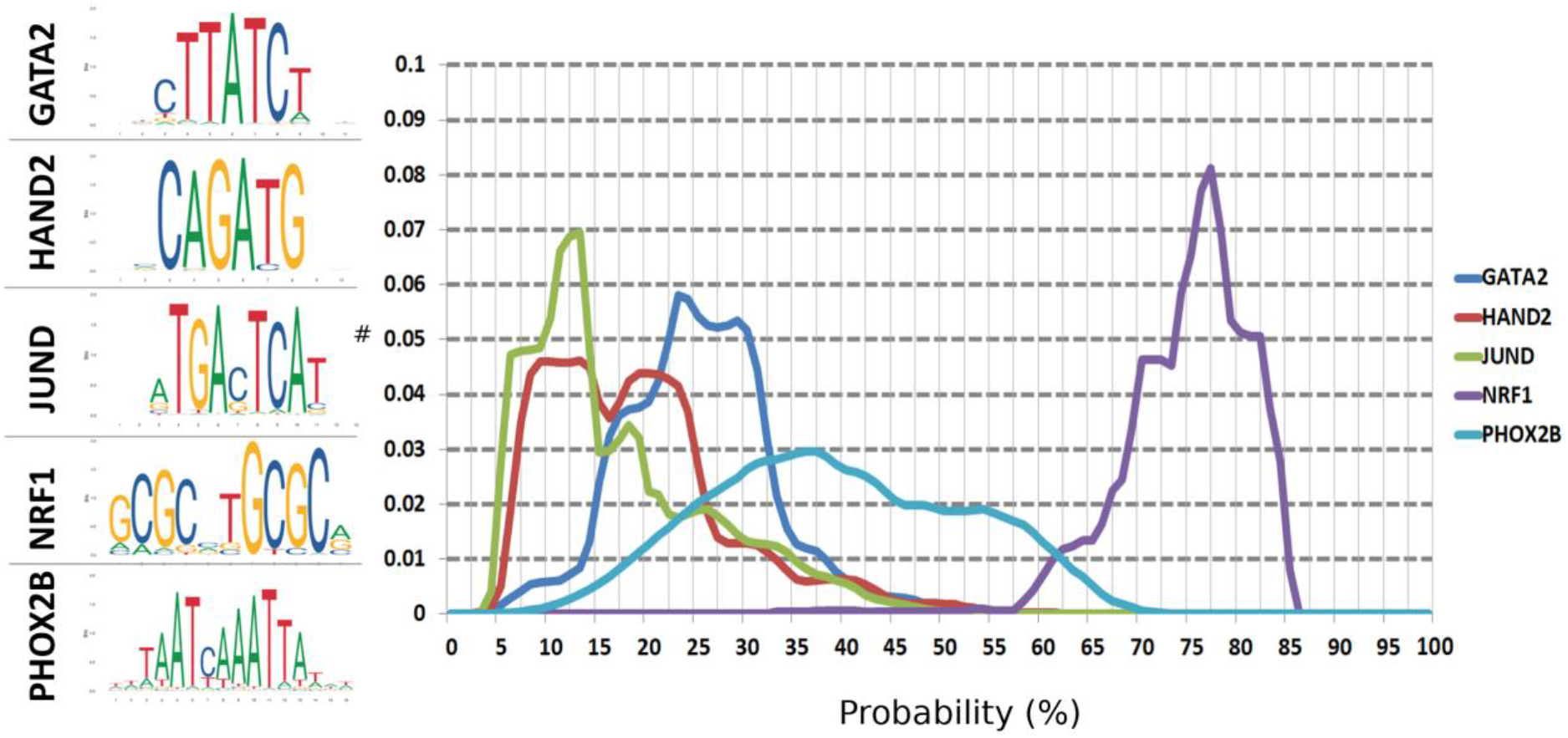
Sequence logos and the distribution of probabilities of detected motifs.

### Conservation of Transcription Factor Binding Sites across different species

The conservation of the TFBS motifs compared to human sequences were also examined individually, for all five transcription factors for each species (Figure 4). At the transcription factor level, it appears that the pattern of occurrences is broadly consistent in the five cases. For NRF1, slightly different conservation pattern becomes visible, however the trends are generally the same for all TFs. For NRF1 less conservation is visible in most cases, with the exception of *Bos taurus* and *Mus musculus*. In general, the conservation closely follows evolutionary patterns, as more closely related species share a higher proportion of motifs with *Homo Sapiens*. The lowest values are present in the case of *Rattus norvegicus* and *Mus musculus,* regardless of the identified TF. These results represent a general trend, but there may be differences at individual motif level

**Figure 4:**
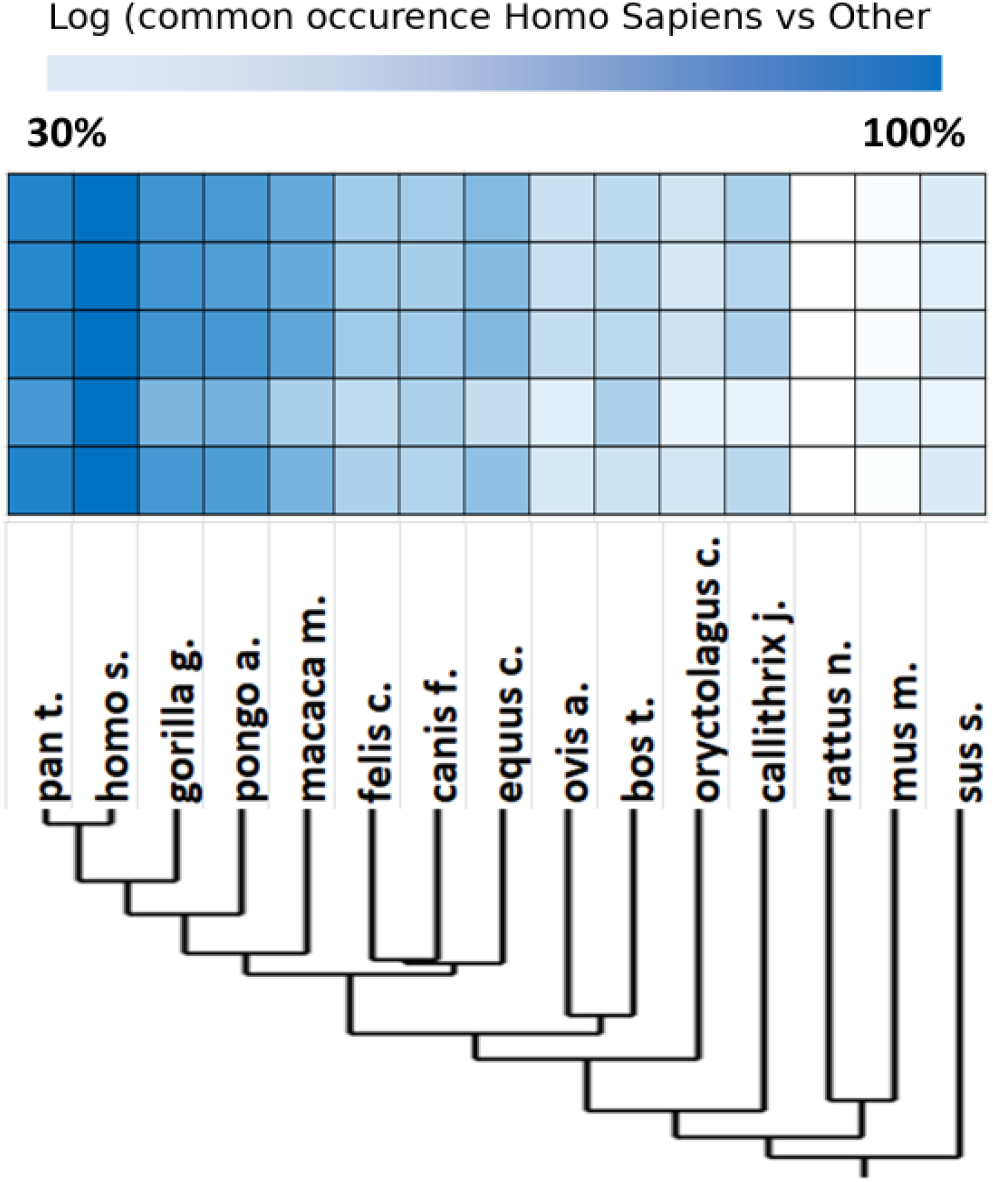
Conservation of motifs across different species and phylogenetic tree calculated from the aligned TFBSs.

### The co-regulation mediates PSD protein interaction network assembly

Regulatory regions affect the transcription of genes, however, their variability (both their distance from regulated regions and in the sequence pattern of the motifs) limits our ability to connect them precisely to their regulated genes. In addition, the transcriptional protein machinery also has a complex composition – multiple regulatory elements and proteins are involved. Therefore, in general, only simplified models can be created for these processes and in our research we also used this approach.

We obtained the coordinates of PSD genes from SynatomeDB and selected those that have a maximum 300 nucleotide distance from the five investigated TFs. In parallel we also used the CHEA3 server to identify co-regulated genes for different TFs (Table 2). Interestingly, NRF1, which has a more specific description but at the same time the least data, has the highest number of co-regulated genes, according to these independent methodologies - this also confirms that the data from NRF1 likely contain the fewest false positives. Next, using the IntAct database we analyzed whether these gene products participate in direct or indirect (a third partner is present that binds to any two proteins from the list) PPIs. Such examples of co-regulated PSD genes by NRF1 participating in the same network are AINX_HUMAN and PPAC_HUMAN, sharing a common interaction partner (KSR1_HUMAN).

**Table 2:** Number of co-regulated genes for each transcription factor, and the number of experimentally verified PPIs between them.

| **Transcription factor** | **Co-regulation (own data)** | **Co-regulation (ChEA3)** | **Direct interaction** | **Indirect interaction** |
| --- | --- | --- | --- | --- |
| JUND | 8 | 6 | 8 | 2 |
| GATA2 | 9 | 7 | 7 | 2 |
| NRF1 | 30 | 23 | 29 | 7 |
| PHOX2B | 15 | 1 | 12 | 2 |
| HAND2 | 20 | 2 | 16 | 0 |

Examining the evolution of regulatory elements may provide possible information from hitherto unknown aspects of the formation of the complex nervous system. Co-regulatory and protein interaction studies may help to explore the complex protein network of PSD, which may also help in the long run to understand neurological diseases. At the time the study was conducted, no single-cell Chip-seq data was available in any public repositories. Therefore we were forced to work with lower resolution data, but our long-term goal would be to perform the highest resolution studies available to access a close up picture of the PSD protein network.

## Acknowledgement

The authors would like to express their greatest gratitude to Laszlo Dobson for his computational help and to Endre Barta for his research advice.

